# Unfolding of RNA via Translocation Through a Nanopore

**DOI:** 10.1101/2021.06.21.449308

**Authors:** Sadhana Chalise, Murugappan Muthukumar

## Abstract

RNA unfolding and refolding are important biological phenomena, which occur during the transfer of genetic information from DNA to RNA to proteins. During these processes, RNA is found in single stranded, secondary and tertiary structures, including secondary conformations like hairpins and pseudoknots. Understanding the diverse conformations of RNA and how these influence the dynamics of unfolding and refolding is crucial to gain insight to fundamental biological processes. In this work, we employ coarse-grained Langevin dynamics simulations of a simple model of different RNA hairpins passing through a geometric nanopore to find the influence of structural changes on the translocation dynamics. The threshold voltage of unfolding depends on the length of the hairpin attached to the tail. The lag time to unfold is longer for smaller applied voltages and for the architectures containing a longer hairpin attached to the tail. Chain translocation dynamics for different architectures are largely collapsed by the threshold. A distinct signature for the base unfolding time was observed for the bases around the unpaired bases in all the RNA hairpin models. These results can motivate future technologies or experiments that use translocation to predict secondary structures of polynucleotides.

## 1 Introduction

Nucleic acids, which constitute some of the biopolymers present in living cells, are essential for the continuity of life because they make up the genetic information of living things.^1,2^ Nucleic acids are found in single stranded, secondary, and tertiary structures such as hairpins and pseudoknots, which play a crucial role in transforming genetic information from DNA to RNA to proteins.

The complex structure of RNA depends on the cellular environment during the transcription,^3,4^ splicing,^5^ translation^6,7^ and protein synthesis.^8^ The genetic information is tran-scripted from DNA to RNA through RNA polymerase. The transcripted information undergoes splicing in which some information is sliced out from the sequence making new sequences. The newly formed sequences are then translated to synthesize one or more proteins. The above processes involves structural changes of secondary and tertiary structures of RNA including at the basic level folding and unfolding events and many also involve the transmission of nucleic acids through protein pores.^3–8^ Therefore understanding how the diverse conformations of nucleic acids fold and unfold and how these complex structures move through nanopores is crucial to gain insight into mechanisms related to the transformation of genetic information from from DNA to RNA to proteins.

Taking inspiration from the biological applications, technologies involving synthetic and biological nanopores have been used in detection of genetic sequences. This technology has been widely applied in DNA sequencing^9–13^ and has been recently applied in RNA and protein sequencing as well.^14–17^ The sequencing of DNA, RNA and proteins is essential in understanding human genomics, gene expression, health care, and the microbiome. Furthermore, such technologies can be applied to determine nucleic acid structures.^15,16,18,19^ With these motivations, the main goal of this project is to understand the role of secondary structures on RNA dynamics during translocation.

With a growing number of studies on DNA, RNA, and protein sequencing, it is very important to understand how the secondary and tertiary structures of RNA folds and unfolds in response to applied external forces. Similarly, it is crucial to study how RNA structure changes during translocation through nanopores. Systematic studies of a single molecule under applied mechanical forces have been done to understand the folding and unfolding mechanisms of complex nucleic acid structures.^11,20–24^ In typical experiments, a constant force is applied using optical tweezers to pull the two ends of DNA/RNA hairpins from which force extension curves are constructed. In these experiments, it is observed that at critical force, RNA hairpins hop between folded and unfolded states.^20,24^ The critical forces are extracted from the force extension curves. Similarly, simulations have been done mimicking these experiments in order to find the folding and unfolding mechanisms of RNA hairpins in response to applied forces for a broad range of temperatures.^21,23^ The hoping between folded and unfolded states of polynucleotide hairpins was observed for hairpins with different stem lengths, loop lengths, and CG content.^11^ In these studies, mechanical force is applied at the two ends of a polynucleotide hairpins, which might be different from the forces relevant to biological processes. However, in translocation, the force is distributed due to interactions between polynucleotides and the pore. Since the interaction of polynucleotides with different protein pores are observed, translocation might provide a better representation of real systems.

A number of studies have been done to understand the unzipping kinetics of polynucleotides during translocation through synthetic and biological nanopores. Translocation is driven by an applied voltage bias, which transmits a force onto the nucleic acid molecule (negative backbone charge), ultimately moving the molecule from one side of the pore to the other. The dependence of unzipping kinetics on pore diameter and threshold voltage have been studied for a wide range of voltages and lengths of hairpins.^25–30^ These studies find that the unfolding process dominates the total translocation time.

Few experiments and little theoretical work has been done focusing on the the influence of the secondary structures of polynucleotides.^31–34^ To obtain the signature of secondary structures of RNA, a new tool combining solid state nanopores and optical tweezers has been proposed to measure the net unfolding force for RNA structures.^32^ However, no systematic studies have been done to understand the role of different architectures on the dynamics of translocation through nanopores.

In this paper, we systematically study the role of hairpin architecture on the unfolding kinetics of RNA during translocation. To initiate studies on this, we employ coarse-grained simulations of a crude model of different RNA hairpins as they translocate through a geometric pore. We restrict to models containing 72 nucleotides of same types of base pairs in five different architectures. Our main focus is to find the influence of structural changes on the translocation dynamics. The results show that the threshold voltage to unfold different RNA architectures is higher when a longer hairpin segment enters the nanopore first. We also study the time evolution of probability of unfolding of all the models for all the applied voltages and extract the lag time as a function of applied voltages for all the architectures. The result show that the lag time is longer for lower applied voltages and for models with longer hairpin segment entering the nanopore first. However, the lag time shows no significant differences for the different models, when collapsed by the threshold voltage. In simulations, we are able to access dynamical information during translocation which is not accessible through experiments, namely base by base unfolding dynamics. The results show that there is a distinct signature of the base unfolding time for bases near the unpaired bases. The signature is due to the lag between the unfolding of the basepair just before a group of unpaired bases and the basepair just after. This gives the foundation for the development of technologies that use translocation to predict secondary structures of polynucleotides.

This article is organized as follows. In Section 2 we describe the model and simulation details that we employ. In Section 3 the results of the simulations are presented, followed by conclusions and a discussion of future directions in Section 4.

## 2 Methods

### Model

The model consists of two parts, an RNA hairpin with a tail and two membrane walls containing a cylindrical pore inside it as shown in Fig. 1. The nucleic acid molecule is represented by a united atom model as described in our previous work.^35^ Each nucleotide contains three spherical beads representing the phosphate, sugar, and base groups, respectively as shown in Fig. 1(c). The phosphate bead carries a charge of -*e*, where *e* is the fundamental unit charge. The beads representing the sugar and base do not carry charge.

**Figure 1:**
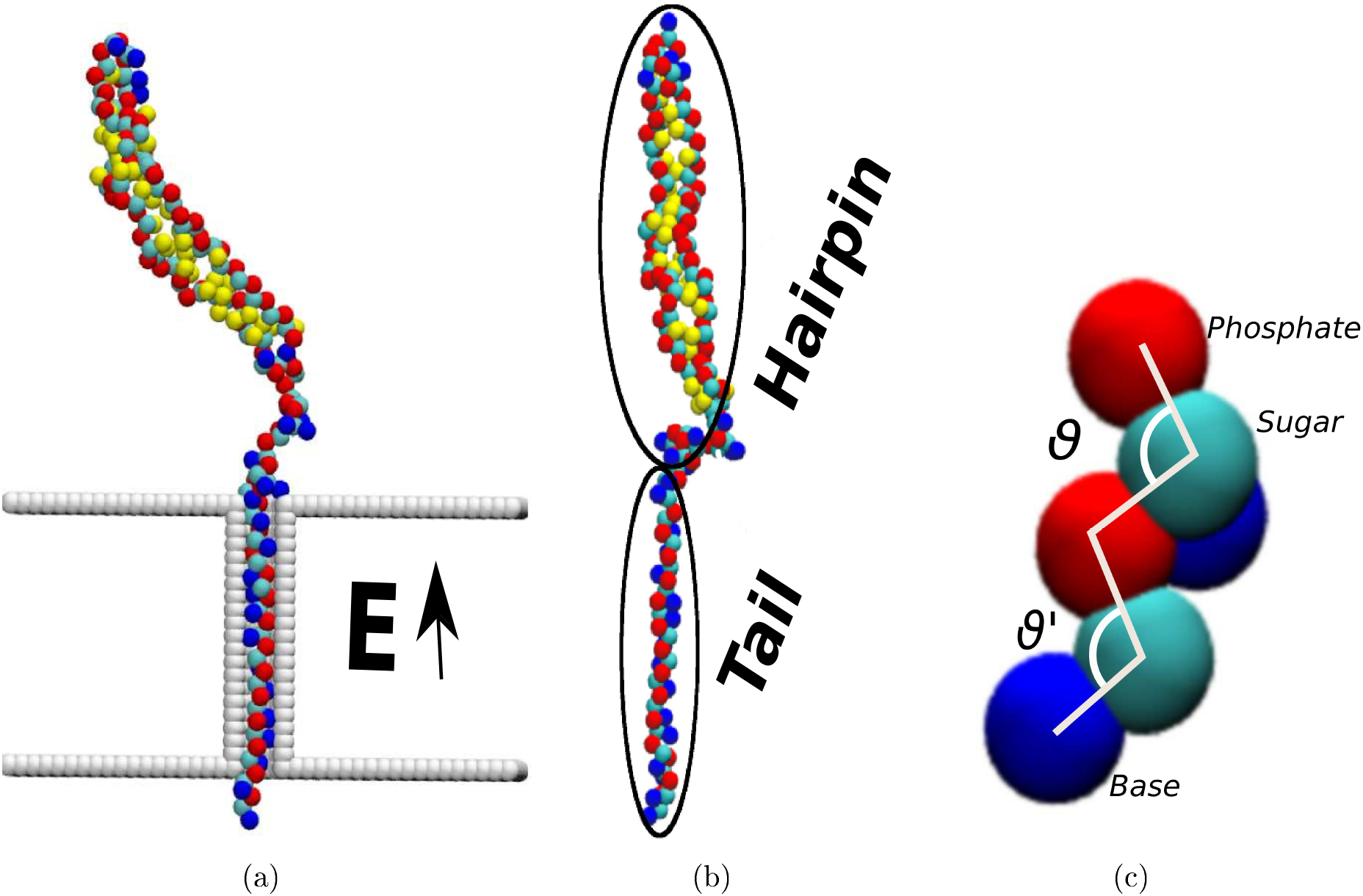
(a) The configuration before starting the simulation; the hairpin remains outside the membrane and the tail is inserted inside the pore. A voltage bias is applied across the membrane generating electric field **E**. (b) The tail and the hairpin section of one model is shown, (c) Representation of three bead unified model, *θ* is a representative backbone angle (phosphate-sugar-phosphate) and *θ*’ is a representative side angle (phosphate-sugar-base), *θ* = *θ*’ in our simulations.

The membrane and the pore walls are represented by spherical beads that do not carry any charge. The stem length of the pore is 50 and the diameter is 14. For simplicity each bead has the same diameter and mass.

We consider five different architectures of the hairpin, each containing 72 nucleotides as shown in Fig. 2. The first model we consider consists of a simple hairpin with 22 base-pairs and a 4 unpaired base loop, also called a tetraloop as represented in Fig. 2 (a) (22HP). The second model consists of an interior loop connecting a stem of 11 base-pairs and a hairpin of 11 base-pairs, which is shown in Fig. 2 (b) (11IL11HP). The third model shown in Fig. 2(c) contains two hairpin domains containing 11 base-pairs in each hairpins and connected by 4 unpaired bases(11-11 HP). The fourth and fifth models are similar to the 11-11 HP model, however, both models contain two hairpins with 16 and 6 base-pairs. In the fourth model, the tail is connected to the 16 base-paired hairpin loop (16-6HP) shown by Fig. 2 (d) and in the fifth model, it is connected to the 6 base-paired hairpin loop (6-16HP) as represented in Fig. 2(e). Each of the hairpin loops has an unpaired tetraloop.

**Figure 2:**
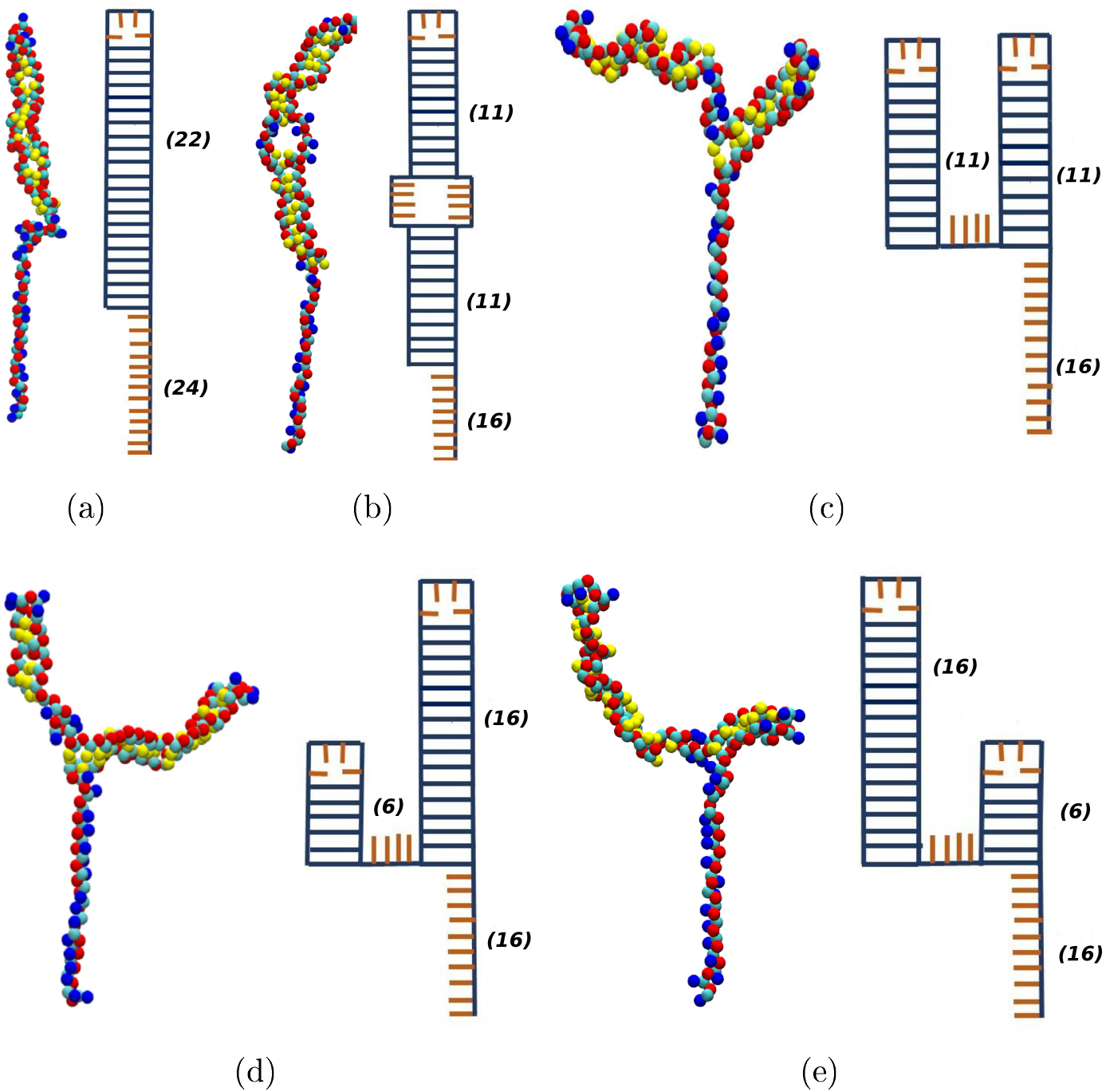
Different architectures of an RNA hairpins with tails. The left image in each frame shows the equilibrated model and the right image shows a 2D structure corresponding to the model. Red beads represent negatively charged phosphate groups, cyan beads represent sugar, yellow beads are paired bases and blue beads are unpaired bases. The models are (a) 22HP, (b) 11IL11HP, (c) 11-11HP, (d) 16-6HP and (e) 6-16HP.

### Simulation details

Langevin dynamics simulations were used to observe the unzipping trajectory of the hairpins under the influence of an applied electric field. The simulation is done using the Large-Scale Atomic/Molecular Massively Parallel Simulator (LAMMPS) package.^36^ The unzipping trajectory is computed by solving the Langevin equation of the *i^th^* bead of the molecule:

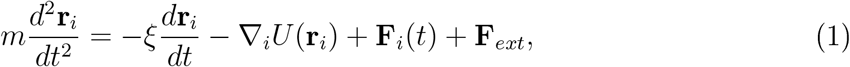

where *m* is the mass of the bead, **r**_*i*_ is the position of the *i^th^* bead, *ξ* is the friction coefficient, *U* is the total interaction potential acting on the *i^th^* bead, **F**_*ext*_ is the force acting on the bead due to the applied external electric field **E**, and **F**_*i*_(*t*) is the random force acting on the *i^th^* bead due to solvent molecules at time t and temperature T. This random force term satisfies the fluctuation dissipation theorem:

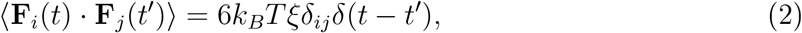

where *k_B_* is Boltzmann’s constant, and *δ*(*t*) is the Dirac delta function.

Simulations were done in dimensionless Lennard-Jones (LJ) units, in agreement with LAMMPS. In defining units, 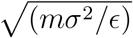 is taken as a unit time and (4*πϵ*_0_*σϵ*)^1/2^ is the unit charge. Each beads has a unit mass, *m* =1 which is equal to 96Da. The beads has diameter of 1 *σ* which is equal to 3.0. The phosphate bead has charge equal to *ϵ* = 1.602 × 10^-19^ C and unit *ϵ* is taken approximately equal to 0.2 *kcal/mol.* For simplicity, the value of the friction coefficient in Eq (1) is choosen arbitrarily to be 1 LJ units. In the simulation, we used an integration time step of *dt* = 0.0002τ LJ units, where, the characteristic time 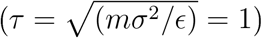 in our simulations.

There are bonded and non-bonded interaction potentials acting on each bead.

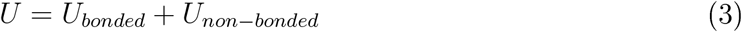

where *U*_bonded_ = *U_b_* + *U_a_* and *U_non-bonded_* = *U_LJ_* + *U_C_* + *U_hb_* are the different potentials acting in the system, which are described below. The bonded potential includes contributions from a spring-like bond (*U_b_*) and an angle potential (*U_a_*). In our system, the bond potential between connected beads (i and i÷l) is represented by a harmonic potential and the angle potential between three successive beads (i, i÷l, and i÷2) is represented by a cosine-squared potential as:

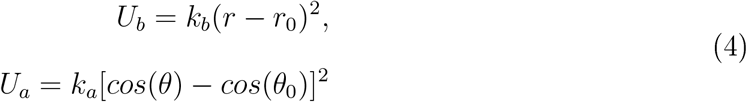

where, *k_b_* = 5344 units (≈ 171 kcal/m ol^2^) is the spring constant,^35^ *k_a_* = 250 units (≈ 60 kcal/mol) is the angle constant,^37^ r is the distance between connected beads, *r*_0_ = 1 is the equilibrium distance, *θ* is angle of backbone and side groups as shown in Fig 1 (c) and *θ*_0_ = 105^0^ is the equilibrium angle.^37^ The values of the parameters are chosen close to realistic values to promote RNA folding.

The non-bonded potential acting on each bead includes three contributions: excluded-volume interactions, a screened Coulomb potential and a Gauss potential to mimic hydrogen bonding.

To model excluded volume interactions, a truncated Lennard Jones (LJ) potential

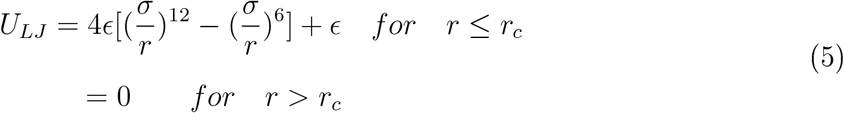

is used where *ϵ* = 1 (≈ 0.2 kcal/mol) is the depth of potential well which is taken as *k_B_T*/3 for temperature *T* = 300 *K* Additionally, *σ* = 3.0 is the effective bead diameter, and the potential is truncated at *r = r_c_* = 1.12*σ*.

The screened Coulomb potential is modeled by using the truncated Debve-Hückel potential as

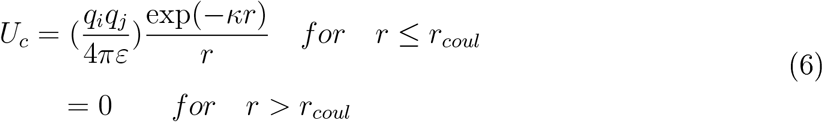

where, *q_i_* denotes the electric charge on bead i, *κ* = 0.639 units is the inverse Debye length for a monovalent salt concentration of 0.6 M, *ε* = *ε*_0_*ε_r_* is the permittivity with *ε*_0_ being permittivity in free space and *ε_r_* being the dielectric const ant, and *r_coul_* = 10/*κ* is the cutoff distance for truncation of the electrostatic potential. Although the effective dielectric constant inside the pore is unknown, our value of the dielectric constant is 80 which is the dielectric constant of water at room temperature. No beads other than the phosphates carry charge. Therefore, the Coulomb potential is only computed between phosphates.

In order to model hydrogen bonding between the bases in the nucleic acid, a Gauss potential,

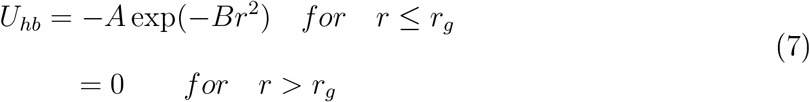

is applied between paired bases where *A* and *B* are parameters with units of energy and distance^-2^, respectively. In RNA, the strength of hydrodgen bonding ranges from 2 to 12 *k_B_T*. For simplicity, we have used A = 10*k_B_T* for the amplitude of the potential. In order to form the desired folded structures we have employed one-to-one interaction, *U_hb_* is applied pairwise between the two specific bases that form a base pair in the desired folded state. Although this model is not realistic, we are trying to understand the first order physics of RNA hairpin unfolding using a crude model.

### Simulation process

To start the simulation, an equilibrated hairpin as shown in Fig. (2) is required. Using a flat initialized configuration, the tail is inserted into the pore and the last bead of the chain was kept fixed. The configuration is allowed to equilibrate for 2000*τ* LJ time units by solving Eq. (1), with no applied field. A Verlet algorithm in LAMMPS^36^ is used to solve the equations of motion. Initially, a random velocity drawn from Boltzmann distribution at a given temperature is also assigned to all moving beads.

The configuration at the end of this equilibration process is taken as the initial conformation for a simulation of unfolding dynamics under the applied electric field. A uniform electric field *E* is applied across the pore. The pore and the membrane are fixed and only the nucleic acid is allowed to move throughout the simulation. Eq. (1) is integrated in time using the Verlet algorithm until the hairpin was either completely unfolded and translocated to the other side or until 5 × 10^7^ time-steps (10, 000*τ* LJ time units) was elapsed. The positions of the beads are recorded after every 1000 time-steps. The diameter of the pore is such that only a single nucleotide is allowed to enter the pore. Therefore, the translocation of the nucleotide is possible only after unfolding has happened. This allows us to study the unfolding dynamics of the RNA hairpin. For a given applied voltage bias, 500 to 1500 runs are performed for each architecture. The number of runs are varied such that we have about 500 successful runs for the analysis of each architecture and applied voltage. For the analysis, only successful unfolding events are taken into consideration.

## 3 Results

In this section, we present the results of our simulations. The results are organized as follows. We first discuss the threshold voltages of unfolding for all the models and the time evolution of probability of unfolding in Subsection A. This is followed by the unfolding dynamics of individual base-pairs in Subsection B.

### 3.1 Fraction of successful unfolding events

For all the models and all applied voltages, we first checked if the hairpin completely unfolded and translocated at the end of the simulation, 10,000τ LJ time units. The ratio of the number of unfolded events to the total number of simulations was calculated and fitted with a sigmoidal function:

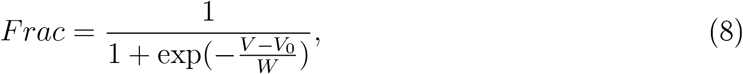

where, *V*_0_ is the voltage at which the inflection point of the Sigmoid curve is observed, and *W* is a measure of the sensitivity of the unfolding fraction to changes in voltage. Fig. (3) shows that the nature of unfolding is similar in all the models where the primary difference is the threshold voltage, defined by *V*_0_-This qualitative behavior isin consistent with single molecule experiment on mechanical pulling of RNA hairpin using optical tweezers.^20^

Table 1 shows the values of fit-constants we obtained. The models 22HP and 11IL11HP show similar threshold voltages. This shows that the internal loop has very little effect on the unfolding mechanism. Among the two domain hairpins (11-11HP, 6-16HP, and 16-6HP), it shows that the threshold voltage depends on the length of the hairpin attached to the tail. When a longer hairpin is attached to the tail, the threshold voltage for unfolding is higher. While the primary differences between the different models is the threshold voltages, the W or sensitivity does slightly vary between the models as well. Table 1 shows that the two domain cases seem to have slightly larger values of W. It is clear that the different structures play a role and further study is needed to understand this.

**Table 1:**
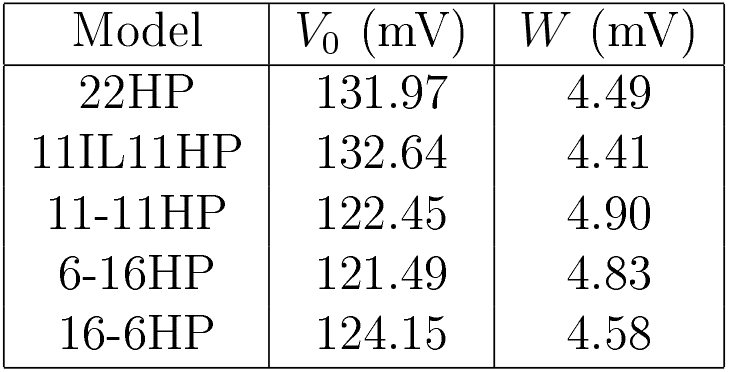
The threshold voltage (*V*_0_) and the *W* of the unfolding fraction for different models, when fit to Equation (8).

Further, we study how the probability of unfolding *(P_unfold_*) evolves with time. We define *P_unfold_* as the ratio of unfolded events to the total number of events at given time for each applied voltage bias and architectures. *P_unfold_* at long time is equal to the fraction of unfolding at different voltages as shown in Fig. 3. Fig. 4 shows how the unfolding of RNA hairpins occurs as a function of time for different architectures and at different applied voltages. Fig. 4 (a) shows the time evolution of *P_unfold_* for the 22HP architecture; it shows that for a given voltage, the unfolding process starts only after a certain lag time. The lag time is shorter for higher voltages. The nature of unfolding of other architectures as shown in Fig. 4 (b-e) is similar to Fig. 4 (a).

**Figure 3:**
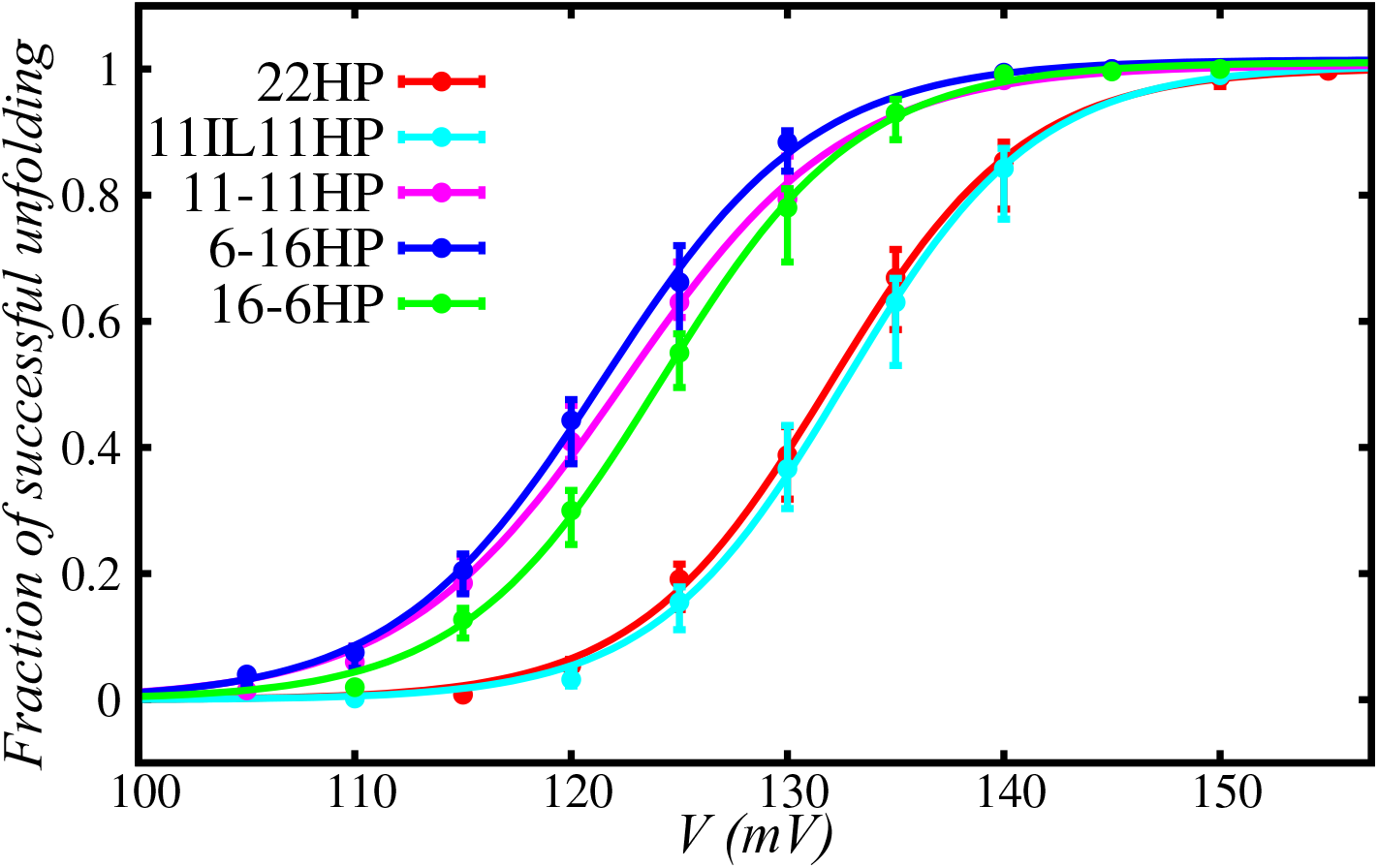
Fraction of successful unfolding events for all models as a function of applied voltage. The curves from top to bottom represent models 6-16HP, 11-11HP, 16-6HP, 22HP and 11IL11HP respectively.

**Figure 4:**
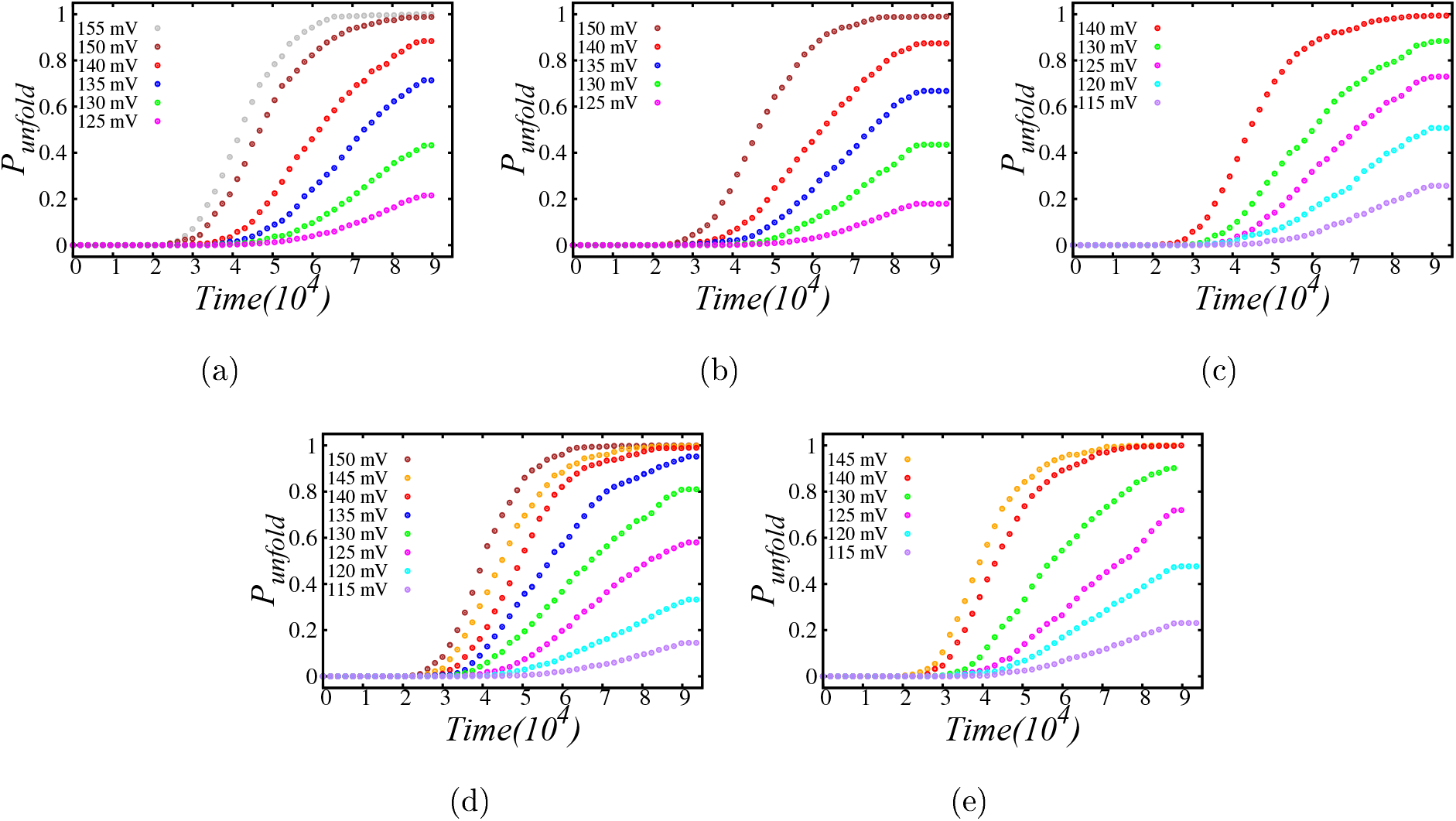
Probability of unfolding as a function of time for models (a) 22HP, (b) 11IL11HP, (c) 11-11HP, (d) 16-6HP and (e) 6-16HP. In all the cases the left-most curve has the highest voltage and the right-most curve has the lowest.

We extracted the lag time from the intersection of the baseline and fitted a line of the evolution of *P_unfold_* as shown in Fig. 5(a). The example is for the case of 155mV applied voltage and 22HP model. Using the same method for all the applied voltages and all the structures, we get the lag time as a function of applied voltages as shown in Fig. 5(b). The red ‘plus’ data points represent the lag time for the 22HP architecture. The figure shows that on increasing the applied voltage, the lag time decreases, as it is easier to unfold the hairpin when more energy is applied to the system. Blue ‘cross’ data points represent the values for the 11IL11HP model. The data points in the curve show that the presence of an internal loop in the model seems to have a minimal effect on the lag time. Other data points show similar behavior for all other architectures but with different time scales. The figure shows that the lag time increases with the increasing length of the hairpin closer to the tail for a constant voltage. Therefore, lag time might be a way to capture the secondary structures of RNA. Moreover, we subtract the threshold voltages of different models as shown in Table 1 from the applied voltages (*V* — *V*_0_) and re-plot the lag time as a function of *V* — *V*_0_ for all the architectures as shown in Fig. 6. The figure shows no significant difference in the lag time between models as a function of *V* – *V*_0_ for all the models. This result implies that the threshold voltage of a model carries the information about the structures of RNA.

**Figure 5:**
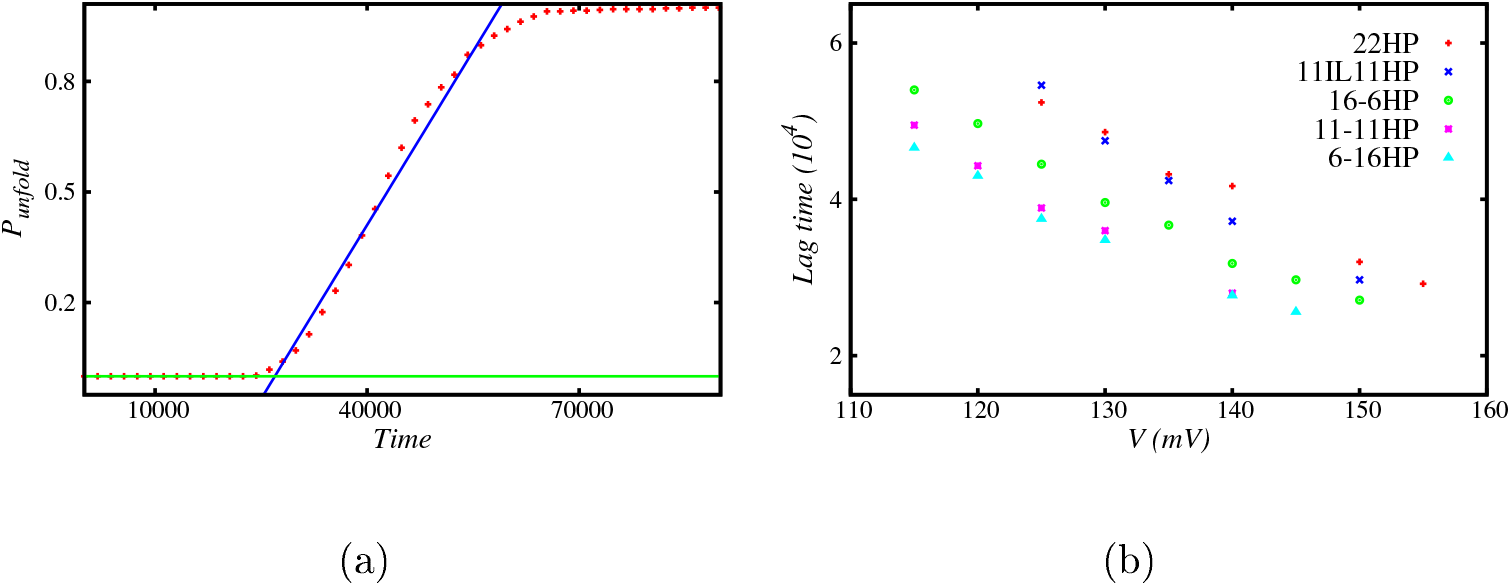
(a) Example of lag time extraction. This is the case of 155 mV for 22HP model, (b) Lag time as a function of applied voltage for different architectures. Lag time is longer for smaller applied voltages and for the structures with longer hairpin attached to the tail.

**Figure 6:**
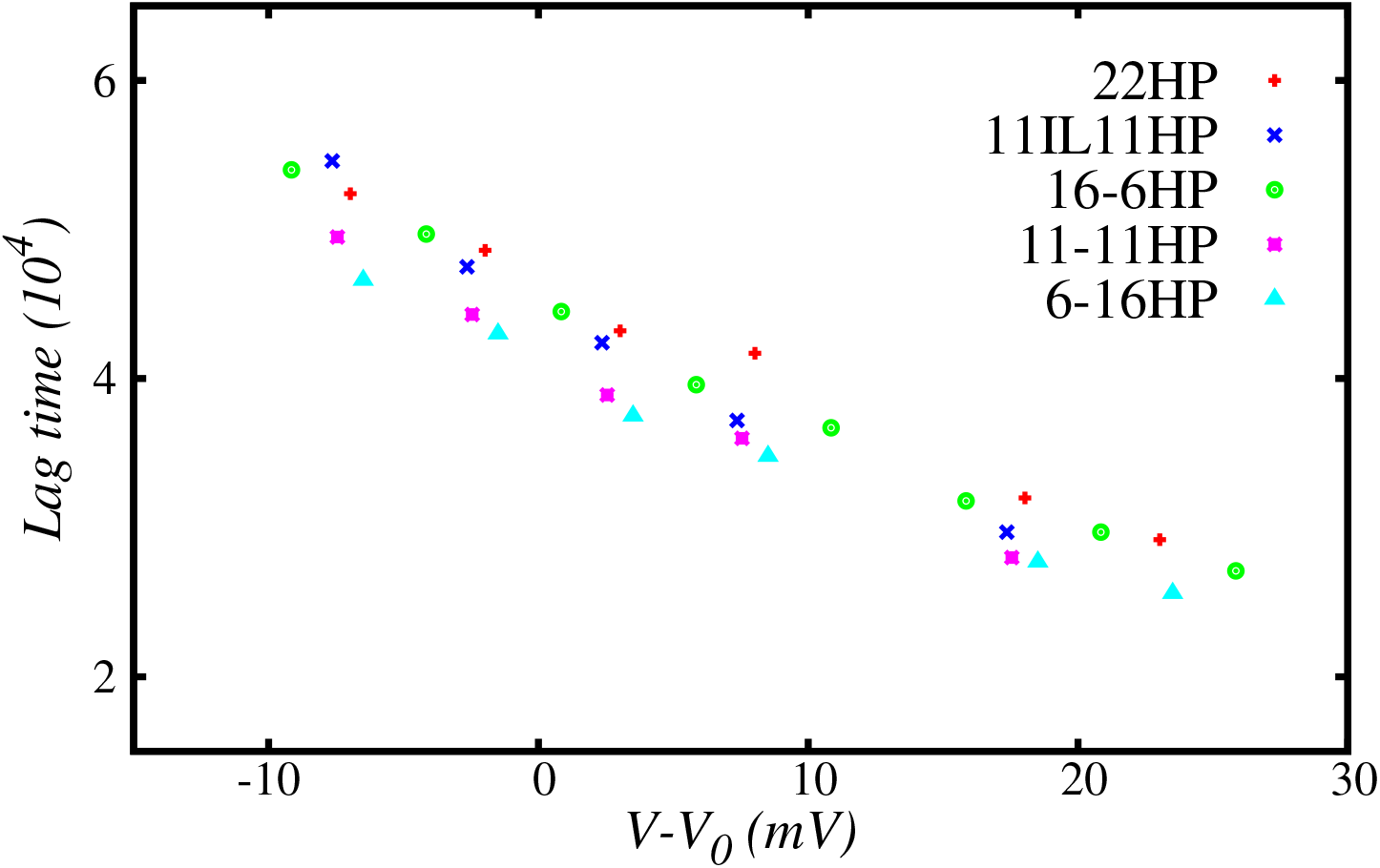
Lag time as a function of difference between applied voltage and threshold voltages for all the architectures. Lag time shows no significant differences for different models.

### 3.2 Unfolding dynamics of individual base-pairs

Here we show how unfolding behavior is observed for each base-pairs in the various models. We define unfolding time of each base-pair as the time it takes a base-pair to permanently unfold; after which it never returns to a folded state. Fig. 7 shows the unfolding time of each base-pair at different applied voltages and for different architectures. Base-pair 1 is the one closest to the tail and base-pair 22 is the one that unfolds the last. From Fig. 7 it is observed that for a hairpin of any length, the unfolding time increases with base-pair number, unfolding occurs sequentially along with the RNA. Additionally, the unfolding is very fast towards the end of the hairpin. We also observed that for two domain architectures (11-11HP, 16-6HP, and 6-16HP), the unfolding of the first few base-pairs of the second hairpin is very fast. After the complete unfolding of the first hairpin loop, the tetraloop can translocate with less force because there is no force (hydrogen bonding in base-pairs) acting against the applied voltage bias. This might have assisted the unfolding of the first few base-pairs of the second hairpin. However, further studies are needed to verify this.

**Figure 7:**
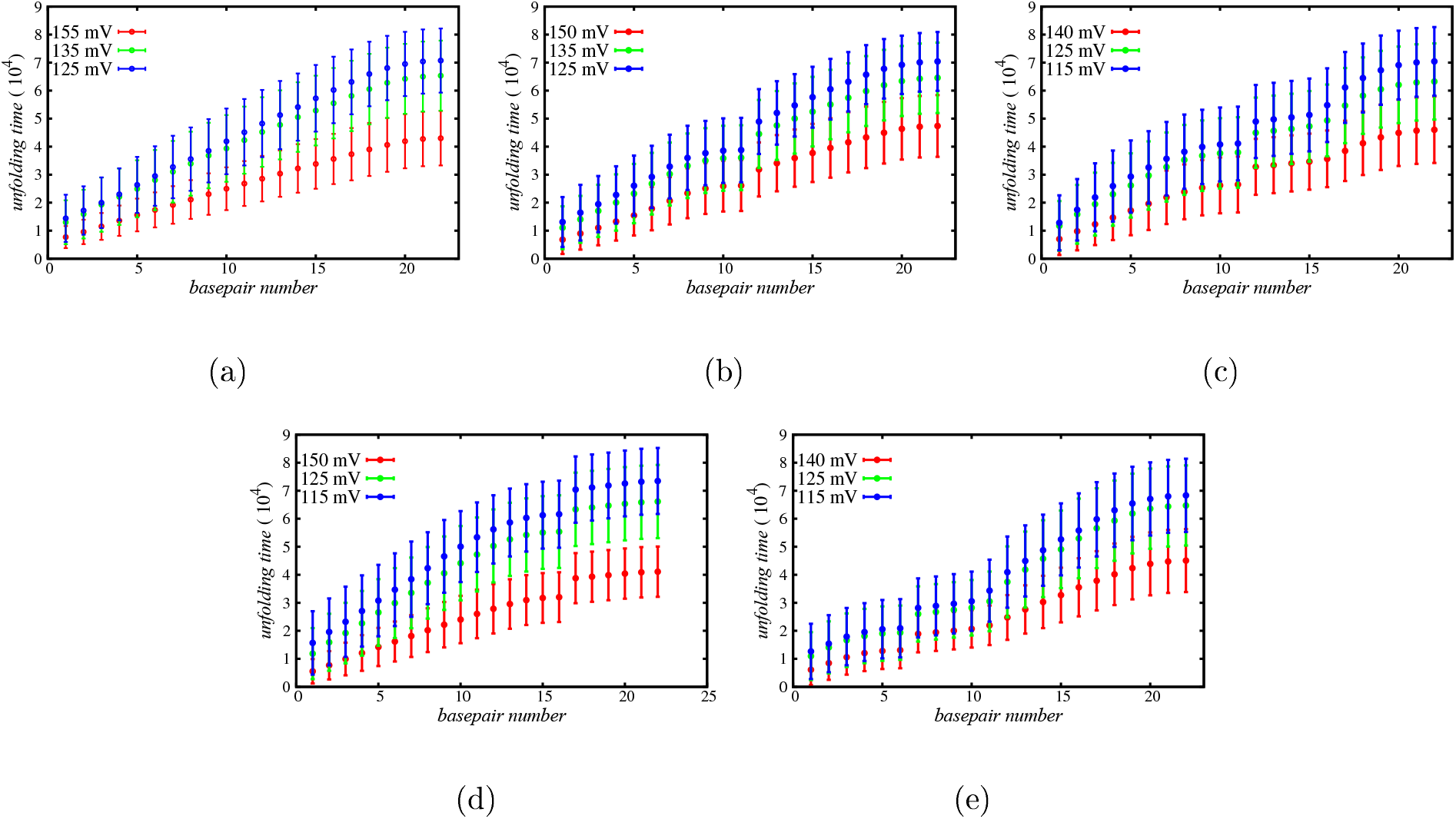
Unfolding time per base-pair for models (a) 22HP, (b) 11IL11HP, (c) 11-11HP, (d) 16-6HP, and (e) 6-16HP. The points are averages from at least 500 simulations and the error bars represent the standard deviations. In all cases the voltages increases moving from the top curve to the bottom one.

Fig. 8 further clarifies this result where we plot the time difference of unfolding of consecutive base-pairs versus base-pair number for different architectures. In Fig. 8(a) (22HP architecture), the first data point shows the time taken to unfold the first base-pair, the second data point shows the time between the unfolding of the first and the second base-pairs, and so on. The result shows that, the time taken to unfold the first base-pair is higher than that for rest of the base-pairs. This is because of the lag time taken by the hairpin to start unfolding. Fig. 8 also shows that, the unfolding of the last few base-pairs is faster compared to other base-pairs. The 11IL11HP architecture shows similar behavior to that of the 22HP architecture, except it shows the unfolding of both the stem and the hairpin domains, as indicated by the two repeated regions as shown in Fig. 8 (b). Fig. 8(c) and (e) are for the 11-11HP and 6-16HP architectures, respectively. Both of them consists of two hairpin domains. The variation of time difference with respect to base-pair numbers of the first hairpin in both cases shows similar behavior to that of the 22HP architecture. The unfolding of base-pairs of the second hairpins however, show qualitatively different behaviors. Once the unfolding of the first hairpin domain is completed, the unfolding of the second hairpin domain begins after a certain lag time. This lag time is due to the existance of the tetraloop in the hairpin. After the unfolding of first base-pair in the second hairpin domain, the unfolding of next few base-pairs is faster and then it slows down for a few base-pairs. Finally, unfolding gets faster for the remaining last few base-pairs. The faster unfolding of the first few base-pairs of the second hairpin domain is due to the assisted force from the tetraloop. Fig. 8 (d) shows results for the 16-6HP architecture. This architecture also consists of two hairpin domains. The variation of time difference with respect to basepair number of the first hairpin shows similar behavior to the rest of the architectures, as explained above. Once the unfolding of the first hairpin is completed, all base-pairs of the second hairpin unfold fast. This is because the hairpin has only six base-pairs.

**Figure 8:**
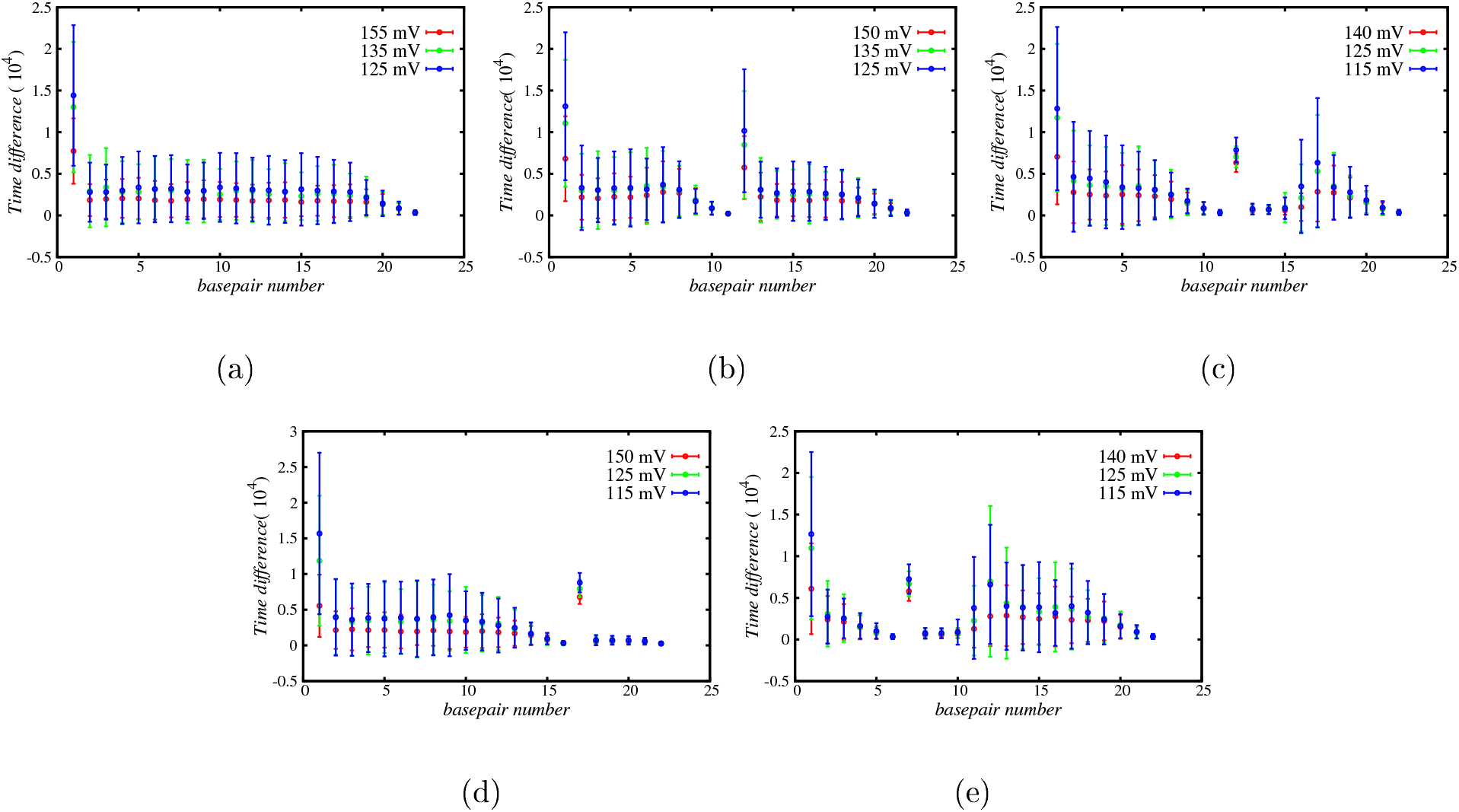
Unfolding time difference per base-pair for models (a) 22HP, (b) 11IL11HP, (c) 11-11HP, (d) 16-6HP, and (e) 6-16HP. The points are averages from at least 500 simulations and the error bars represent the standard deviations. Results for all voltages overlap within the error bars.

## 4 Conclusions

In this study we have employed a simple model of various RNA hairpins and Langevin dynamics simulations to study the unfolding of RNA. We have used the united atom model to model nucleotides and a simple cylindrical pore to model the nanopore. We have considered five different architectures of the hairpin, each containing 72 nucleotides and 22 base-pairs. The first model is 22HP representing a simple hairpin with 22 base-pairs. The second model is 11IL11HP, made up of a stem of 11 base-pairs connected to a hairpin of 11 base-pairs by an interior loop, third, fourth, and fifth models are 11-11HP, 16-6HP, and 6-16HP respectively. They consists of two hairpin domains connected by four unpaired bases. The numbers in the name of the model are the numbers of base-pairs in each hairpin domains. Each of the hairpin domain has an unpaired tetraloop.

The results of this study provide the foundation for development of technology that uses translocation to predict secondary structures of polynucleotides. The threshold voltage of unfolding depends on the length of the hairpin attached to the tail. The longer the hairpin attached to the tail, the higher the threshold voltage. In addition, the lag time to unfold is longer for lower applied voltages and for the models with longer hairpins attached to the tail. However, the lag time collapses when the voltage is adjusted by the threshold voltage. Moreover, the base by base unfolding dynamics show that there is a distinct signature of base unfolding time for the bases before and after the unpaired bases in all the RNA hairpin models.

In the future, more complex models of secondary and tertiary structures of polynucleotides and different geometries of pores can be used to understand the role of RNA architectures on the dynamics of translocation through nanopores and to predict the secondary and tertiary structures of polynucleotides.

## Funding

Acknowledgement is made to the National Institutes of Health (Grant No. 5R01HG002776-16), the National Science Foundation (DMR-2015935), and the AFOSR Grant FA9550-20-1-0142 for financial support.

## Notes

### Competing Interest Statement

The authors have declared no competing interest.

## References

(1) Blanco, A.; Blanco, G. In Medical Biochemistry; Blanco, A., Blanco, G., Eds.; Academic Press, 2017; pp 121–140.

(2) van Holde, K. E.; Zlatanova, J. In The Evolution of Molecular Biology; van Holde, K. E., Zlatanova, J., Eds.; Academic Press, 2018; pp 87–94.

(3) Pan, T.; Sosnick, T. RNA folding during transcription. Annu. Rev. Biophys. Biomol. Struct. 2006, 35, 161–175.

(4) Pedersen, J. S.; Bejerano, G.; Siepel, A.; Rosenbloom, K.; Lindblad-Toh, K.; Lander, E. S.; Kent, J.; Miller, W.; Haussler, D. Identification and classification of conserved RNA secondary structures in the human genome. PLoS Comput Biol 2006, 2, e33.

(5) Warf, M. B.; Berglund, J. A. Role of RNA structure in regulating pre-mRNA splicing. Trends in biochemical sciences 2010, 35, 169–178.

(6) Kozak, M. Regulation of translation via mRNA structure in prokaryotes and eukaryotes. Gene 2005, 361, 13–37.

(7) Ramakrishnan, V. Ribosome structure and the mechanism of translation. Cell 2002, 108, 557–572.

(8) Svoboda, P.; Cara, A. D. Hairpin RNA: a secondary structure of primary importance. Cellular and Molecular Life Sciences CMLS 2006, 63, 901–908.

(9) Wanunu, M. Nanopores: A journey towards DNA sequencing. Physics of life reviews 2012, 9, 125–158.

(10) Heng, J. B.; Ho, C.; Kim, T.; Timp, R.; Aksimentiev, A.; Grinkova, Y. V.; Sligar, S.; Schulten, K.; Timp, G. Sizing DNA using a nanometer-diameter pore. Biophysical journal 2004, 87, 2905–2911.

(11) Woodside, M. T.; Behnke-Parks, W. M.; Larizadeh, K.; Travers, K.; Herschlag, D.; Block, S. M. Nanomechanical measurements of the sequence-dependent folding landscapes of single nucleic acid hairpins. Proceedings of the National Academy of Sciences 2006, 103, 6190–6195.

(12) Venkatesan, B. M.; Bashir, R. Nanopore sensors for nucleic acid analysis. Nature nanotechnology 2011, 6, 615–624.

(13) Manrao, E. A.; Derrington, I. M.; Laszlo, A. H.; Langford, K. W.; Hopper, M. K.; Gillgren, N.; Pavlenok, M.; Niederweis, M.; Gundlach, J. H. Reading DNA at singlenucleotide resolution with a mutant MspA nanopore and phi29 DNA polymerase. Nature biotechnology 2012, 30, 349–353.

(14) Smith, A. M.; Jain, M.; Mulroney, L.; Garalde, D. R.; Akeson, M. Reading canonical and modified nucleobases in 16S ribosomal RNA using nanopore native RNA sequencing. PloS one 2019, If, eO2l67O9.

(15) Soneson, C.; Yao, Y.; Bratus-Neuenschwander, A.; Patrignani, A.; Robinson, M. D.; Hussain, S. A comprehensive examination of Nanopore native RNA sequencing for characterization of complex transcriptomes. Nature communications 2019, 10, 1–14.

(16) Xie, W.; Chipman, J. G.; Robertson, D. L.; Erikson, R.; Simmons, D. L. Expression of a mitogen-responsive gene encoding prostaglandin synthase is regulated by mRNA splicing. Proceedings of the National Academy of Sciences 1991, 88, 2692–2696.

(17) Depledge, D. P.; Srinivas, K. P.; Sadaoka, T.; Bready, D.; Mori, Y.; Placantonakis, D. G.; Mohr, L; Wilson, A. C. Direct RNA sequencing on nanopore arrays redefines the transcriptional complexity of a viral pathogen. Nature communications 2019, 10, 1–13.

(18) Lockhart, D. J.; Winzeler, E. A. Genomics, gene expression and DNA arrays. Nature 2000, 405, 827–836.

(19) Kono, N.; Arakawa, K. Nanopore sequencing: review of potential applications in functional genomics. Development, growth & differentiation 2019, 61, 316–326.

(20) Liphardt, J.; Onoa, B.; Smith, S. B.; Tinoco, I.; Bustamante, C. Reversible unfolding of single RNA molecules by mechanical force. Science 2001, 292, 733–737.

(21) Hyeon, C.; Thirumalai, D. Mechanical unfolding of RNA hairpins. Proceedings of the National Academy of Sciences 2005, 102, 6789–6794.

(22) Hyeon, C.; Thirumalai, D. Forced-unfolding and force-quench refolding of RNA hairpins. Biophysical journal 2006, 90, 3410–3427.

(23) Liu, F.; Ou-Yang, Z.-c. Monte Carlo simulation for single RNA unfolding by force. Biophysical journal 2005, 88, 76–84.

(24) Harlepp, S.; Marchal, T.; Robert, J.; Leger, J.; Xayaphoummine, A.; Isambert, H.; Chatenay, D. Probing complex RNA structures by mechanical force. The European Physical Journal E 2003, 12, 605–615.

(25) Zhao, Q.; Comer, J.; Dimitrov, V.; Yemenicioglu, S.; Aksimentiev, A.; Timp, G. Stretching and unzipping nucleic acid hairpins using a synthetic nanopore. Nucleic Acids Research 2008, 36, 1532–1541.

(26) Mathé, J.; Visram, H.; Viasnoff, V.; Rabin, Y.; Moller. A. Nanopore unzipping of individual DNA hairpin molecules. Biophysical Journal 2004, 87, 3205–3212.

(27) McNally, B.; Wanunu, M.; Meller. A. Electromechanical unzipping of individual DNA molecules using synthetic sub-2 nm pores. Nano letters 2008, 8, 3418–3422.

(28) Dudko, O. K.; Mathé, J.; Szabo, A.; Moller. A.; Hummer, G. Extracting kinetics from single-molecule force spectroscopy: nanopore unzipping of DNA hairpins. Biophysical journal 2007, 92, 4188–4195.

(29) Sauer-Budge, A. F.; Nyamwanda, J. A.; Lubensky, D. K.; Branton, D. Unzipping kinetics of double-stranded DNA in a nanopore. Physical Review Letters 2003, 90, 238101.

(30) Stachiewicz, A.; Molski, A. Diffusive dynamics of DNA unzipping in a nanopore. Journal of computational chemistry 2016, 37, 467–476.

(31) Wang, X.; Li, Y.; Li, T.; Liu, L.; Wu, H.-C. The effect of secondary structures on the generation of characteristic events during the translocation of DNA hybrid through α-hemolysin. Science China Chemistry 2016, 59, 135–141.

(32) van den Hout, M.; Vilfan, I. D.; Hage, S.; Dekker, N. H. Direct force measurements on double-stranded RNA in solid-state nanopores. Nano letters 2010, 10, 701–707.

(33) Gerland, U.; Bundschuh, R.; Hwa, T. Translocation of structured polynucleotides through nanopores. Physical biology 2004, 1, 19.

(34) Schink, S.; Renner, S.; Alim, K.; Arnaut, V.; Simmel, F. C.; Gerland, U. Quantitative analysis of the nanopore translocation dynamics of simple structured polynucleotides. Biophysical journal 2012, 102, 85–95.

(35) Muthukumar, M.; Kong, C. Simulation of polymer translocation through protein channels. Proceedings of the National Academy of Sciences 2006, 103, 5273–5278.

(36) Plimpton, S. Fast parallel algorithms for short-range molecular dynamics. Journal of computational physics 1995, 117, 1–19.

(37) Klimov, D.; Betancourt, M.; Thirumalai, D. Virtual atom representation of hydrogen bonds in minimal off-lattice models of *a* helices: effect on stability, cooperativity and kinetics. Folding and Design 1998, 3, 481–496.

